# Cell-free DNA from ascites is a suitable source for the identification of driver mutations and aligns well with functional chemoresistance testing in late-stage ovarian cancer patients

**DOI:** 10.64898/2026.11.22.768385

**Authors:** David Holthaus, Mahdi Rasa, Christoph Rogmans, Huy Duc Le, Annika Brauer, Mouhamad Khouja, Monika Brüggemann, Franziska Kaiser, Kalle Günther, Katrin Rodenkirchen, Uwe Oelmüller, Ibrahim Alkatout, Nicolai Maass, Andre Franke, Burkhard Brandt, Jörg-Paul Weimer, Marion T. van Mackelenbergh, Dirk O. Bauerschlag, Michael Forster, Nina Hedemann

## Abstract

**Aim:** Ovarian cancer is the leading cause of cancer-related deaths among gynaecological malignancies. This is mainly caused by late diagnosis and resistance development to standard-of-care chemotherapy consisting of platinum compounds and taxanes. Most patients develop resistance to therapy in the first 2 years, leading to high rates of recurrence. In these patients, surgical therapeutic options are limited, and so is the possibility of resampling of tumour material for precision medicine purposes. However, almost 100% of the patients with recurrences accumulate malignant ascites that is filled with tumour components such as single tumour cells, tumour spheroids, and cell-free DNA. Malignant ascites therefore provides a straightforward, minimally invasive means of resampling recurrent ovarian cancer for (functional) precision medicine.

**Methods:** We sampled ascites of a cohort of 22 late-stage ovarian cancer patients and sequenced cell-free DNA using a custom-made targeted sequencing panel. In parallel, 25 spheroid lines of 17 patients and 12 organoid lines of 9 patients were successfully established and subsequently sequenced using the same panel. Identified molecular vulnerabilities like variants in the insulin-growth factor pathway and PIK3CA were then tested on patient-derived spheroids using a titration of standard-of-care chemotherapy in comparison with small-molecule inhibitors targeting these pathways.

**Results and Conclusion:** Sequencing of cell-free DNA in ascites was applicable for longitudinal tracking of driver mutations as well as identification of driver mutations in single samples. Variants in ascites and cell-culture models aligned well, with organoids showing higher retention of variants identified in ascites. However, the spheroid model was fully sufficient for targeting driver mutations *in vitro* and aligned well with targetable molecular vulnerabilities.

We thus propose that cfDNA from ascites is a suitable source for the identification of driver mutations that aligns well with functional chemoresistance testing in late-stage ovarian cancer patients.

## Background

Ovarian cancer has the highest mortality rate among gynaecologic tumours ^1,2^. High-grade serous ovarian carcinomas (HGSOCs) account for 70-80% of the mortality ^2^. Due to the lack of early symptoms, in almost 75% of cases, the diagnosis of ovarian cancer is made only at an advanced disease stage ^3^. This is associated with limited treatment options and reduced overall survival. In addition, up to 80% of patients develop a relapse within the first 2 years after diagnosis, 60% of patients even within the first 6 months ^2^. One of the main reasons is the development of resistance to therapeutic agents, which results in tumour recurrence and metastasis.

The accumulation of ascites is a common phenomenon in ovarian cancer, especially in HGSOC. While at diagnosis only around 30% of patients present with ascites, this number grows to almost 100% in later stages and relapses ^4-6^. The presence of malignant ascites is associated with poorer progression-free survival (PFS) and overall survival (OS) in ovarian cancer ^5,7^. Moreover, volume and frequency of ascites accumulation are associated with poorer outcome ^4,5^. As first-line therapy almost always consists of primary debulking surgery followed by combinatorial chemotherapy of carboplatin with taxanes, malignant ascites is often associated with resistant tumours as it appears later in disease progression after adjuvant chemotherapy ^8^. However, second cytoreductive surgery is uncommon, so sampling of tumours is complicated ^5,8^. Others have proposed the sampling of ascites fluid and its associated cellular content for the prediction of therapeutic success ^9,10^. Werner, et al. ^10^ underline the suitability of ascites for the identification of targetable mutations and to map tumour evolution. However, functional testing was not performed. The use of ascites-derived cell-free DNA (cfDNA) in combination with the use of cellular ascites contents for functional precision medicine purposes would therefore enable targeted therapies of individual mutational profiles and altered pathways ^11^.

To investigate treatment responses and to generate functional resistance data to chemotherapeutics, the primary, patient-derived culture of resected tumour material is necessary. These so-called patient avatars reflect individual patient characteristics, like mutational load and associated resistance patterns. The suitability of two-dimensional (2D) monolayer cell culture models has been debated in this context ^12^ as these models do not reflect the *in vivo* situation of most tumours ^13,14^. Therefore, many studies have shifted from traditional monolayer studies to more complex three-dimensional (3D) systems. One of these is the 3D tumour spheroid model. In tumour spheroids, the cancer cells have a 3D architecture and show oxygen, nutrient and pH gradients, which correspond to the most important features of tumour biology in solid tumours ^12,15^. Tumour spheroid models are comparatively low-cost alternatives to traditional cell culture as they can be grown in off-the-shelf media and ultra-low attachment plates. A second model used for the generation of patient avatars is the organoid model that has been continuously expanded across entities since its first description. Cancer-derived lines have also been established for HGSOC tissue^16-18^.

In this study, we sampled ascites from 22 late-stage ovarian cancer patients and sequenced cfDNA using a custom-made targeted sequencing panel. In parallel, 25 spheroid lines from 17 patients and 12 organoid lines from 9 patients were successfully established and subsequently sequenced using the same panel. Cell culture models showed high concordance to cfDNA from ascites and driver mutations were retained in most patients. Moreover, functional testing showed that these variants could also be targeted *in vitro*.

## Material and methods

### Extraction of cells from ascites

Primary cells were obtained from late-stage ovarian cancer patients undergoing routine surgery (Ethical approval 488/23 & 631/24) at the Department of Gynaecology and Obstetrics in Kiel between 2018 and 2025. Primary cells from ascites were isolated following established protocols ^19,20^. In short, ascites from ovarian cancer patients was washed with DPBS and spun down at 300 x g at 4°C for 5 min. If high numbers of erythrocytes were present in the ascites, erythrocyte lysis was performed using the RBC lysis buffer kit (BioLegend) following the manufacturer’s guidelines. Following erythrocyte lysis, cells were again washed with DPBS and pelleted at 300 x g at 4°C for 5 min.

### Cell culture, spheroid conversion, and harvest

For spheroid cultures, cells were directly transferred to tissue culture flasks with RPMI-1640 medium (PanBiotech) containing 10% foetal bovine serum (Gibco Life Technologies), 1% L-Glutamine (Sigma-Aldrich) and 1% penicillin-streptomycin (Biochrom). After sufficient growth, cells were trypsinized with 0.25% Trypsin-EDTA (Gibco) for 5 min at 37°C, pelleted at 300 x g at 20°C for 7 min, counted using a CellDrop BF (DeNovix) automated cell counter, and seeded into BIOFLOAT 96-well cell culture plates or Corning 96-well Spheroid Microplates (#4520) for spheroid formation. After three days, spheroids were collected, pelleted at 300 x g at 20°C for 7 min, and lysed with genomic lysis buffer supplemented with 0.1% β-mercaptoethanol of the Quick-DNA Miniprep kit (Zymo).

### Organoid establishment and harvest

For organoid cultures, cells from ascites were directly seeded in Matrigel following published protocols ^21^. After polymerization, Matrigel domes were overlaid with ovarian cancer medium 1 (OCM1), OCM1+, and OCM2 (**Supplementary table 1** for medium conditions) with an approximately comparable number of wells per medium condition. Cells were monitored for growth for up to six weeks, until growth was observed and the medium with the best growth condition could be chosen. After establishment, cells were regularly passaged as described before ^22,23^. For harvest of DNA, organoids from regular routine cultures below the 8^th^ passage were used.

For epithelial cells derived from healthy fallopian tubes (FT), cultures were established following published procedures ^22,24^. Cultures were established from patients undergoing routine surgical procedures. Patients with pathogenic conditions of the female reproductive tract, such as malignancies or endometriosis, were not considered for cell isolation. Fallopian tube cells were propagated in FT expansion medium (**Supplementary table 1**) as described for the generation of fallopian tube organoids. Cells were directly seeded into spheroid plates after two-dimensional culture ^22^.

For the harvest of nucleic acids, the Matrigel inside the medium in each well was disrupted using a P1000 pipette tip by scraping of the droplet inside of the medium and collected in a 0.1 % BSA-coated 15 mL tube. The tube was filled up with ice-cold DPBS and organoids were pelleted by centrifugation for 5 minutes at 300× g, 4 °C. After aspiration of the supernatant, 1 mL of cell recovery solution (Gibco) per 4 wells of organoids was added and the mixture was incubated on ice at 4 °C for 60-120 min for the removal of Matrigel. After the incubation period, organoids were again washed with ice-cold DPBS, pelleted by centrifugation for 5 minutes at 300× g, 4 °C, and lysed with RLT buffer supplemented with 2 mM DTT of the AllPrep DNA/RNA kit (QIAGEN).

### Functional testing of chemoresistance

The protocols were adapted from previous studies ^19,23,25^. 3,000 primary ovarian cancer cells were seeded in ultra-low attachment (ULA), black-transparent 96-well U-bottom plates (Corning, 4520) to form spheroids. After 24 hours of growth, cells were incubated with 0.5x CellTox Green cytotoxicity dye (Promega, G8741), and combined with either Cisplatin (obtained from the Clinical Pharmacy Services, UKSH, Campus Kiel), Alpelisib, or Linsitinib (both Cayman Chemical) in concentrations ranging from 0.1-20 µM. Live Cell Imaging measurements were performed using the CELLAVISTA 4 automated cell imager in combination with the SYBOT X-1000 with CYTOMAT 2 C-LiN system (all SYNENTEC). Wells were automatically imaged every 24 hours for a total of 48 hours and afterwards fluorescence data and images were analysed and subsequently extracted with the YT-Software (SYNENTEC) using the Spheroid Count (2F) application. The following settings were used: Exciter: Blue (475/28) - Emission filter: Green Filter 530nm (530/43). In short, the application detects the spheroids in the brightfield image, and analyses the average fluorescence intensity (background-corrected) within this spheroid mask.

After 48 hours, the CellTiter-Glo 3D Cell Viability Assay was performed following manufacturer’s instructions. In short, excess supernatant from the plates was removed and the reagent was mixed 1:1 with PBS and pipetted onto the wells. Well contents were mixed and allowed to stabilize at RT for 60 min. Afterwards, luminescence was detected using a Sparks plate reader (Tecan).

### DNA extraction

DNA was extracted using the Quick-DNA Miniprep kit (Zymo) for spheroids and the AllPrep DNA/RNA Mini kit (Qiagen, #80204) for organoids following the manufacturer’s guidelines. For cfDNA isolation, 1mL of ascites supernatant was thawed and 3mL PBS was added. Automated cfDNA isolation was performed on the QIAsymphony SP (QIAGEN # 9001297), using the QIAsymphony DSP Circulating DNA Kit (QIAGEN #937556) in combination with the 4ml circDNA_4000_DSP Protocol (QIAsymphony DSP Circulating DNA Kit Instructions for Use (Handbook) 06/2022) in accordance with the kit manufacturer’s instructions. The ascites cfDNA was stored at -80°C. For buffy coat gDNA isolation, 200µl of the separated buffy coat sample were processed using the automated QIAsymphony DSP DNA Mini Kit (QIAGEN #937236) with the DNA_Buffy_Coat_200_V7 DSP protocol with 200µl elution volume (QIAsymphony® DSP DNA Kit Instructions for Use (Handbook) Version2) in accordance with the kit manufacturer’s instructions.

### Library preparation and sequencing

The ascites cfDNA eluates and peripheral blood buffy coat gDNA eluates were thawed and used for library preparation using the QIAseq Targeted DNA Pro library preparation Kit for Illumina (QIAGEN #333675) and QIAseq Targeted DNA Pro UDI Set A (96) (QIAGEN #333455) with a customized Ovarian Cancer Panel (GeneGlobe ID: CPHS-52721Z-6415) in accordance with the kit manufacturer’s instructions (QIAseq Targeted DNA Pro Handbook 12/2022). The target enrichment was performed using PCR with a single locus-specific primer per target. The library method included a sequencing adapter with a random unique molecular identifier (UMI) of 12 nucleotides length that was copied to the template sequence in the first PCR. Subsequent amplification was performed in PCR cycles using a universal primer that binds to the adapter sequence. The UMIs were intended for bioinformatic mitigation of sequencing errors stemming from the PCR and the sequencing platform. The libraries were sequenced on NovaSeq 6000 S4-lane (Illumina) using 2×150bp paired end reads, with 12GB of sequencing output planned per sample.

### Bioinformatic analysis

#### Variant filtering and data processing

The QIAGEN CLC Workbench software was configured to identify the UMI sequences in the FASTQ reads, trim the reads, align the sequences to human reference hg38, perform UMI-based error correction and generate bam and bai files containing the consensus reads that could be further analysed. The UMI-based error correction used read group sizes of at least 2 reads, a consensus threshold of 0.51, and performed trimming of non-consensus read ends. Minimum consensus read length after trimming was 20, otherwise consensus reads were discarded. The bam files were imported into GenSearchNGS, and sensitive raw variant calling was performed with 0% VAF threshold and 1 consensus read threshold for a variant call. The raw calls were annotated and filtered with the following GenSearchNGS filtering thresholds: >1 consensus read, type: worse than synonymous, population minor allele frequency (MAF) < 0.001, at least 1 forward and 1 reverse consensus read supporting the variant call. The filtered variant calls and their annotation for all libraries were exported as a single matrix in csv format.

Targeted sequencing variant calls were further processed in R ^26^. Variant tables were imported using the *data*.*table* package (v1.17.8) ^27^, and sample metadata were imported from Microsoft Excel using *readxl* ^28^. Data processing and reshaping were performed using the *dplyr, tidyr*, and *stringr* packages from the *tidyverse* ecosystem ^29,30^. Technical quality filtering retained variants detected in at least one sample with an AF ≥5%, sequencing coverage ≥60×, and ≥5 supporting variant reads.

To reduce recurrent technical artefacts and background variants, two additional filtering steps were performed. First, variants detected at AF >5% in healthy control samples were removed. Second, background variability was assessed using HC and buffy coat samples. For each variant, the background AF range was defined as the difference between the 95th and 5th percentile VAF values in background samples, whereas the biological AF range was calculated as the difference between the maximum and minimum AF values across non-background samples. Variants were retained only if the biological VAF range was at least twofold greater than the corresponding background range or if no background measurements were available. The resulting filtered dataset was used for all downstream analyses.

#### Targeted sequencing data analysis

Samples were assigned to predefined biological groups (cell-free ascites, buffy coat, spheroids, organoids, healthy controls) based on metadata annotations. Data processing and reshaping were performed using the *data*.*table, dplyr, tidyr, stringr*, and *readxl* packages ^28-30^. To identify germline variants, variants detected in matched buffy coat samples with a VAF ≥20% were annotated as germline for the corresponding patient.

A cutoff of 2.5% VAF was chosen for all subsequent analyses. Variant burden was calculated as the number of detected variants per sample and summarized across biological sample groups. VAF distributions were visualized at both the individual-sample and biological-group levels. Single-nucleotide variants were classified into the six canonical pyrimidine-based substitution classes (C>A, C>G, C>T, T>A, T>C, and T>G), while insertions and deletions were grouped as INDELs. Mutational spectra were summarized by biological group and individual sample.

For gene-level analyses, variants were assigned to all annotated genes after removal of non-canonical identifiers (e.g., Ensembl gene identifiers, mitochondrial genes, microRNAs, long non-coding RNAs, and antisense transcripts). VAF heatmaps were generated using *ComplexHeatmap*^31^, with rows grouped by gene and columns ordered according to patient metadata. Germline variants were retained in visualization panels and indicated by an asterisk. Oncoprint-like plots were constructed from the 50 most frequently mutated genes, with samples ranked according to total targeted mutation burden and genes ordered by recurrence across samples. Only genes mutated in more than one patient were plotted.

#### Patient-matched ascites-culture comparisons

To evaluate variant retention during cell culture establishment, patient-matched comparisons between ascites cfDNA and corresponding spheroid or organoid cultures were performed. Only patients with both an ascites cfDNA sample and the corresponding culture type were included. For each matched comparison, variants were classified as *Shared, Ascites-only*, or *Culture-only*. Variant counts were summarized across all matched patients separately for spheroids and organoids and visualized as stacked bar plots.

### Basic statistical analysis and visualization

Basic calculations were performed using MS EXCEL 2021 (Microsoft). IC_50_ determinations were performed in Prism (GraphPad) using non-weighted least squares non-linear regression in the *‘[inhibitor] vs normalized response* – *Variable slope’* setting. Figures and subfigures were plotted using Prism (v11.0.0, GraphPad) and RStudio (v2026.01.0). The panel composition and annotations of the main figures were created using Affinity Designer (v2.6.5, Serif). Schematics were assembled in Affinity and at BioRender.com.

## Results

### Cohort and study design

Twenty-two patients with late-stage ovarian cancer were included in the study (**Figure 1A,B**), 18 of which were pathologically diagnosed as high-grade serous ovarian cancer (HGSOC), 2 as low-grade serous ovarian cancer (LGSOC), 1 as cancer of unknown primary (CUP), and one as granulosa cell tumour (Figure 1C). Median age was 62 years (Range: 25-78) (**Figure 1C**). Median pre-operative CA125 (Cancer Antigen 125/MUC16) was 432 (Range: 29-3540, **Figure 1C**) which was not significantly correlated to PFS (p = 0.689). Nine patients achieved R0 status, while 13 patients were scored R1 after surgery. Sixteen tumours were classified as FIGO stage III, five as stage IV, and one as stage II.

**Figure 1:**
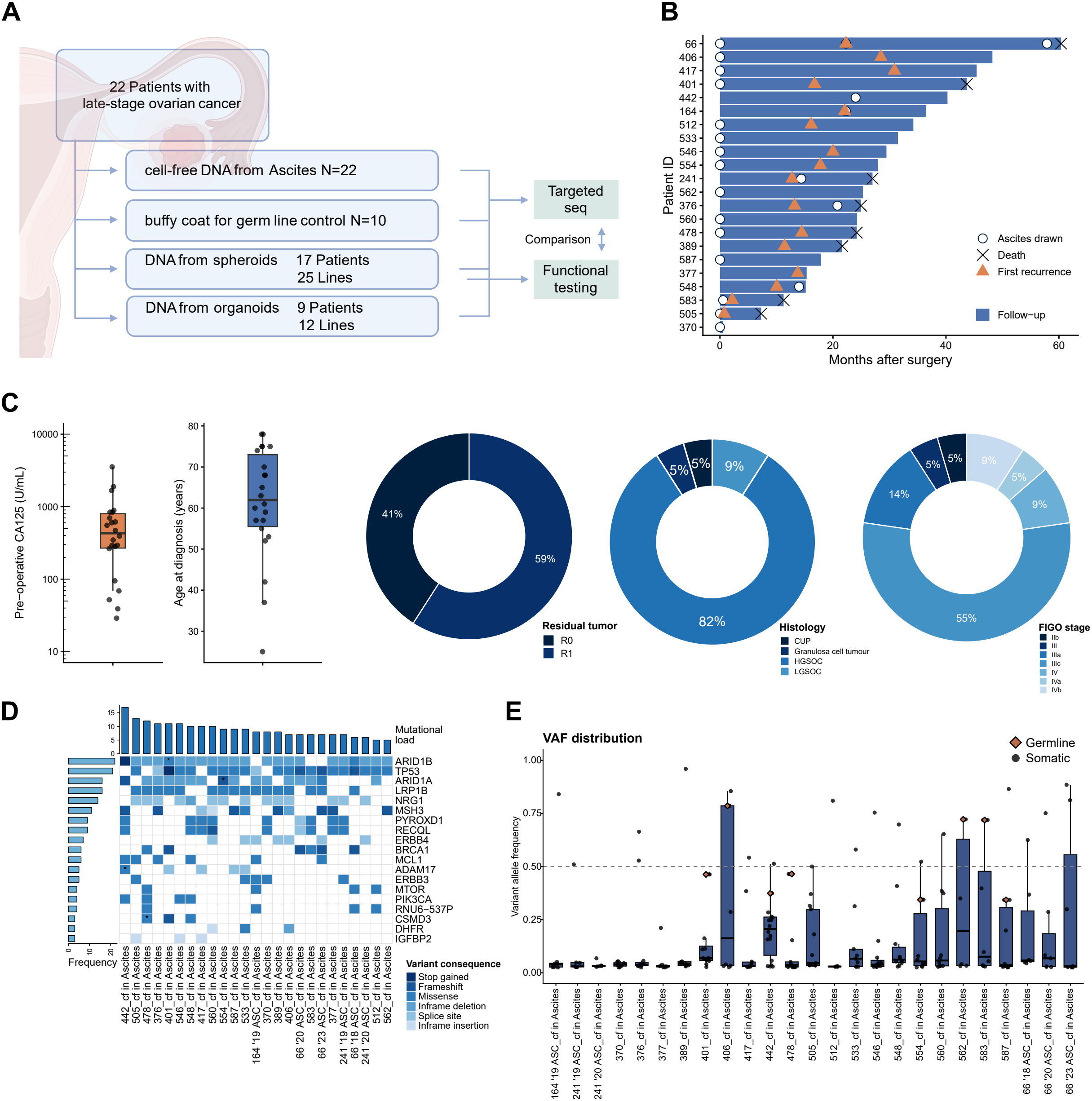
Cohort description and targeted sequencing of cfDNA from late-stage ovarian cancer patients. **A** Schematic representation of the study. **B** Swimmer plot of all included patients. **C** Pre-operative CA125, age at diagnosis, resection status, tumour histology, and FIGO staging of the included patients. **D** Oncoprint-like plot showing mutational load, frequency of mutated genes, and variant consequence of sequenced cfDNA isolated from ascites. **E** Variant burden and **F** variant allele frequency distribution per sample.

cfDNA from ascites was collected from all patients and subsequently sequenced using a custom-made targeted panel of common driver mutations such as *tumour protein p53* (TP53), *BRCA1/2 DNA Repair Associated* (BRCA1/2), *phosphatidylinositol-4,5-bisphosphate 3-kinase, catalytic subunit alpha* (PIK3CA), and *MutS Homolog 3* (MSH3), and other genes commonly associated with resistance such as *insulin-like growth factor 1 receptor* (IGF1R) and *insulin-like growth factor-binding protein 2/5* (IGFBP2/5) of the *insulin-growth factor* (IGF) pathway. Cells from ascites were isolated and cultured in cell culture flasks for spheroid generation and in ECM for organoid generation. In parallel, tumour-derived cell lines were cultured whenever tumour tissue was extracted during surgery for comparison with ascites derived cells. 25 spheroid lines of 17 patients and 11 organoid lines of 9 patients were successfully established and were sequenced using the same targeted panel (**Figure 1A**).

### Sequencing of cfDNA in ascites shows known driver mutations and patient heterogeneity

Sequencing of cfDNA in ascites revealed highest mutational loads in *AT-Rich Interaction Domain 1A/B* (ARID1A/B), TP53, *LDL Receptor Related Protein 1B* (LRP1B), *Neuregulin 1* (NRG1) and *MutS Homolog 3* (MSH3) as expected for an ovarian cancer cohort (**Figure 1D**). Germline mutations were found in three patients in *CUB And Sushi Multiple Domains 3* (CSMD3), ARID1A and B, respectively. Variant burden ranged from 6 to 17 variants per patient (**Figure 1D**). Variant allele frequency (VAF) distributions varied between individuals with most patients showing VAFs of < 10% while others like UF406, UF442, and UF562 showed heterozygous and homozygous pathogenic variants (**Figure 1E**).

### Cell culture models show concordance with variants identified in cfDNA from ascites

Longitudinal sampling of cfDNA during disease progression showed intra-patient specific mutational shifts and changes in variant VAFs (**Figure 2A**). Loss of variants was mainly observed in low-VAF mutations, such as g.56093923T>G in ERBB3 in UF241. Next, we investigated whether mutations detected in ascites are retained in cell culture models. Plotting variants as oncoprint-like plots we could confirm that mutational load was dominated by cell culture samples such as for UF442, UF478, and UF583 while ARID1A/B, TP53, LRP1B, NRG1 and MSH3 dominated by frequency (**Figure 2B**). Spheroids and organoids showed variation in VAFs, but mainly carried similar mutations (**Figure 2C**). Cell culture models derived of ascites were similar to those derived from tumour tissue in their detected variants and VAFs (**Figure 2D**).

**Figure 2:**
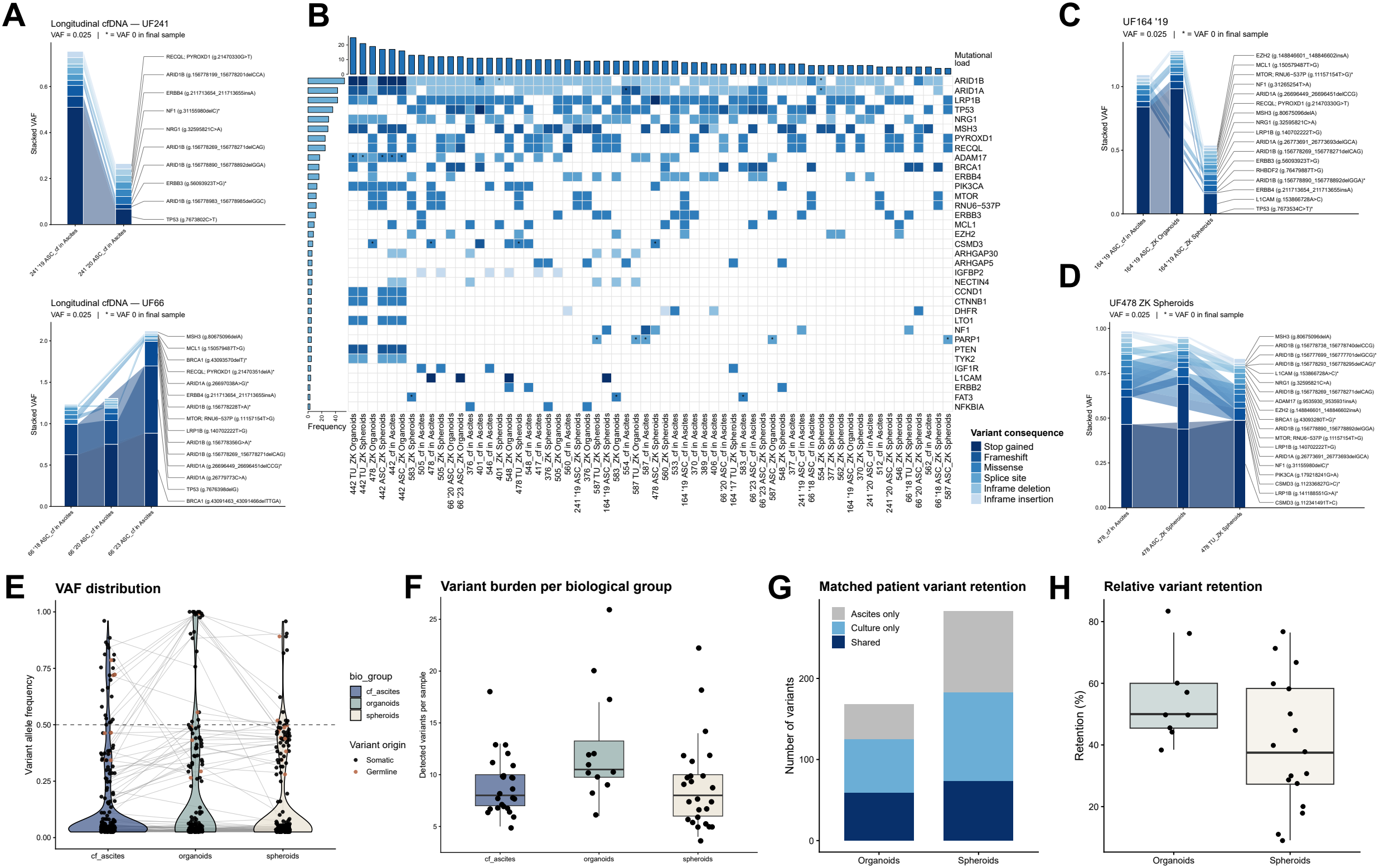
Overlap and retention of pathogenic variants identified in cfDNA from ascites to cell culture models. **A** Longitudinal sampling of ascites of two different patients shows retention of driver mutations with varying VAFs. **B** Oncoprint-like plot showing mutational load, frequency of mutated genes, and variant consequence of cfDNA isolated from ascites, spheroids and organoids. Asterisks indicate germline mutations. **C** Comparison of spheroids and organoids derived from the same ascites sample. Plot shows the retention of most mutations but with lower VAFs in spheroids. D Comparison of ascites and tumour-derived models of the same patients shows high concordance of driver mutations in the cell culture models. **E** VAF distribution of variants present in cfDNA from ascites, spheroids, and organoid cultures. Grey lines link individual mutations per patient identified in more than one group. Red dots mark germline variants while somatic variants are marked in black. **F** Comparison of variant burden per group. **G** Total number of retained variants identified in organoids and spheroids. Variants only found in cfDNA from ascites, shared variants and variants only found in cell culture conditions are shown. **H** Relative variant retention per organoid and spheroid line in comparison to the respective cell-free ascites sample is shown. A cutoff of >0.025 VAF was chosen for all plots.

When comparing overall VAF distribution of cfDNA in ascites versus that of cell culture models, VAFs in ascites were dominated by variants with lower frequencies and only some accumulations below 0.5 and 1 VAF indicating heterozygous and homozygous variants (**Figure 2E**). In organoids and spheroids, the variants with higher frequencies increased. Heterozygous variants were increased especially in spheroids, while homozygous variants were increased in organoids, indicating selection during culture (**Figure 2E**). Comparison of the variant burden per model showed similar results for spheroids and cfDNA from ascites, while the burden in organoids was elevated (**Figure 2F**). When comparing variant retention between models by total numbers, both spheroids and organoids showed comparable numbers of shared mutations to cfDNA samples (**Figure 2G**). At the same time relative variant retention showed higher retention in the organoid model, with a median of 50% compared with 37.5% for spheroids (**Figure 2H**). A comparison of the characteristics of the three groups is given in **Supplementary figure 1**.

When comparing VAF and mutational overlap accounting for germline mutations, the models showed retention of variants (**Figure 3**). Germline mutations such as splice site variant g.226361545_226361546insGG in Poly [ADP-ribose] polymerase 1 (PARP-1) were retained in all models of UF587. It is notable that the spheroid model appeared to select against TP53 mutations, while the organoid models were able to retain these mutations in the cell culture models (**Figure 3**). Some genes appeared to show multiple new low VAF variants such as triple nucleotide deletions in ARID1A/B. The analysis also revealed targetable variants in genes such as BRCA1/2, *beta-catenin* (CTNNB1), PIK3CA, TP53, and the IGF pathway members IGF1R and IGFBP2.

**Figure 3:**
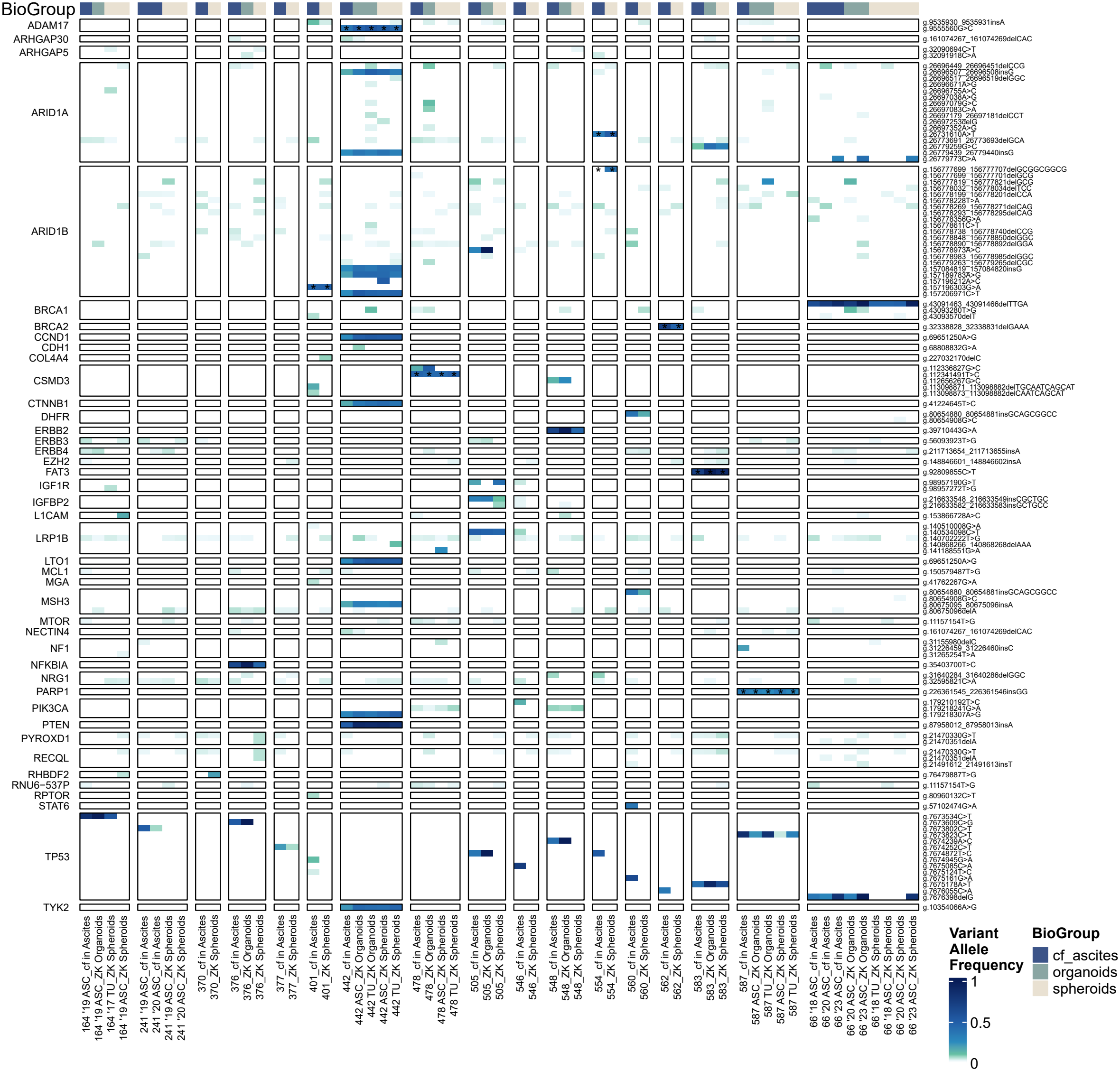
Heatmap showing the variant allele frequency of all matched ascites-cell culture pairs. Heatmap borders indicate individual patients and genes. Individual variants are annotated on the right side. Asterisks indicate germline mutations.

### Functional testing of ascites-derived spheroid cultures corresponds to identified mutational drivers

After identifying mutational profiles and retention of driver mutations from ascites to cell culture models, we assessed whether targeting of these mutations *in vitro* results in corresponding sensitivity profiles. For this, we established a multiplex workflow that tracks effects of chemotherapy on cytotoxicity and viability of ovarian cancer-derived spheroids (**Figure 4A**). Sequencing identified two hotspots of targetable mutations in patients UF442 and UF505 (**Figure 3, Figure 4B**). UF442 showed a g.179218307A>G missense mutation (rs397517201) in PIK3CA while UF505 showed mutations in the IGF pathway with a g.98957190G>T missense mutation (rs746049391) in IGF1R and a g.216633548_216633549insCGCTGC in-frame insertion in IGFBP2. We continued to culture ascites-derived cells from these patients in ultra-low attachment plates to form spheroids. We then treated these spheroids after 24 hours of aggregation time with increasing concentrations of Cisplatin, PI3K-pathway inhibitor Alpelisib, and IGF1R-pathway inhibitor Linsitinib for 48 hours (**Figure 4C,D**) and assessed cytotoxicity by live cell imaging. As a comparison, we additionally included spheroids from patients UF560, UF554, and UF562, as well as two lines of healthy primary fallopian tube epithelium (UF609 and UF613). Imaging revealed high cytotoxicity of Cisplatin and Alpelisib in UF442, intermediate to low cytotoxicity in UF560 for all tested compounds, and high cytotoxicity of Cisplatin and Linsitinib in UF505 (**Figure 4C,D**). Additionally, we assessed viability of the cultures as end-point readout and calculated IC_50_s using dose-response curves. All tested cultures showed high resistance to Cisplatin with IC_50_s ranging from 14.58 to 39.32 µM except for UF442 with an IC_50_ of 2.96 µM. For Alpelisib, UF442 again showed the lowest IC_50_ with 0.48 µM while the other lines showed IC_50_s ranging from 2.553 to 14.97 µM. Targeting IGF1R with Linsitinib resulted in IC_50_s ranging from 3.061 to 16.38 µM with UF505 having the lowest IC_50_ (**Figure 4E,F**). The functional testing supported the findings of the sequencing analysis by confirming vulnerabilities in PIK3CA in UF442 and the IGF pathway in UF505 (**Figure 4F**).

**Figure 4:**
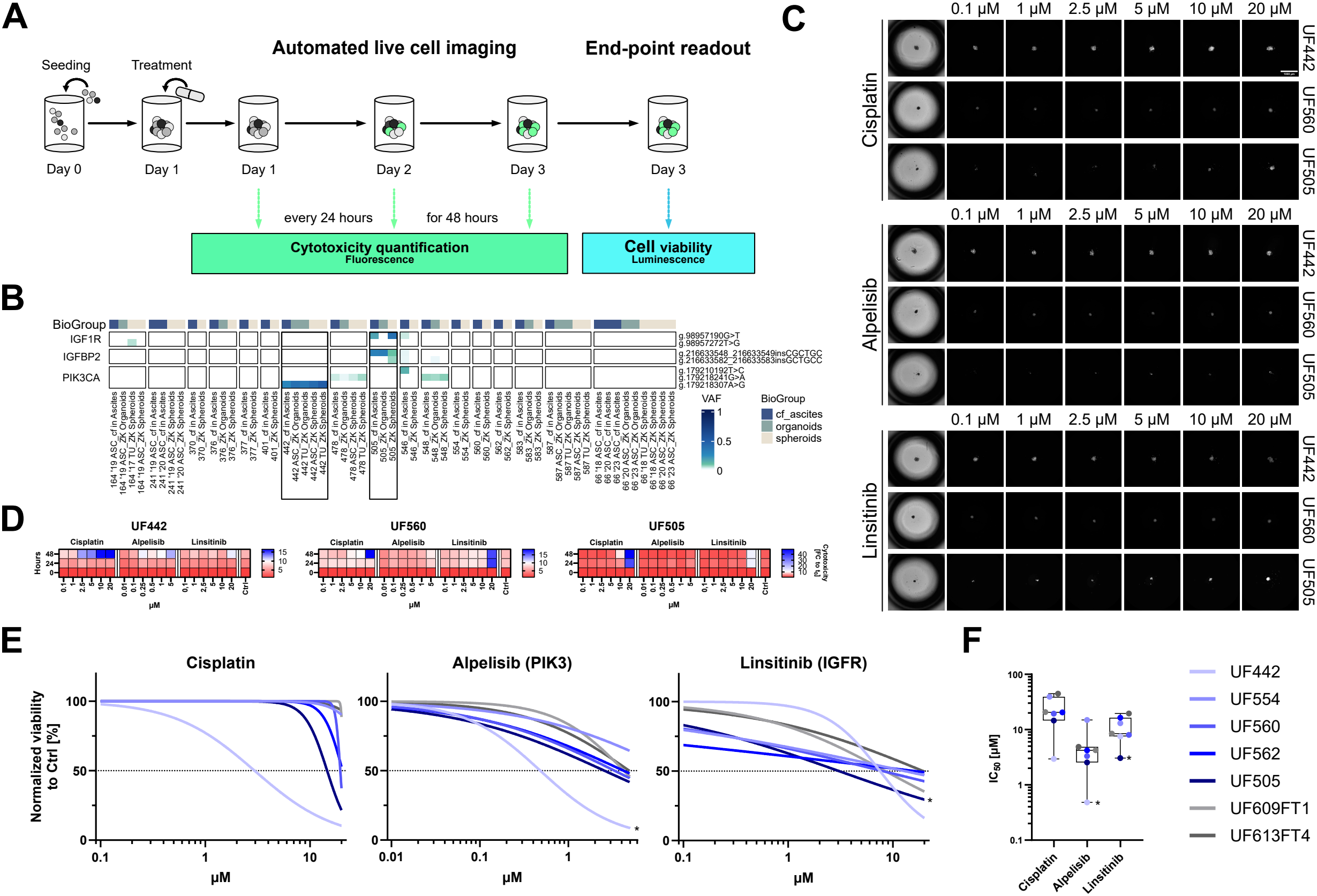
Functional testing of ascites-derived spheroids from ovarian cancer confirms vulnerabilities identified during targeted sequencing. **A** Schematics of the functional testing workflow. Ascites-derived cells are seeded into ultra-low attachment plates, allowed to form spheroids for 24 hours, and incubated with chemotherapy for 48 hours. Live cell imaging is performed during this incubation period followed by a cell viability end-point readout. **B** Zoom in on Figure 3 for the identification of targetable pathways PIK3CA in UF442 and the IGF pathway in UF505. **C** Representative images of the spheroids after 48 hours incubation with the indicated concentration of chemotherapy. Fluorescence of CellTox Green dye indicating cytotoxicity is shown in false colours. White indicates sensitivity. **D** Quantification of the fluorescence over the time course of 48 hours. Means of three independent experiments per line are shown. **E** Endpoint readout using the Cell Titer Glo 3D assay normalized to untreated spheroids. Plots show fitted curves calculated on results of three independent experiments per line. **F** Boxplots of IC_50_s calculated from E. Asterisks indicate the line carrying mutations in the targeted pathway in **E** and **F**.

## Discussion

There is an unmet clinical need for the sampling and targeting of driver mutations in ovarian cancer using (functional) precision medicine. Malignant ascites has been proposed as a medium for sampling as it provides an opportunity for tracking tumour progression and evolution ^5^. New developments in sequencing technologies allow for the isolation and sequencing of cfDNA. cfDNA sampling provides for unbiased spatial sampling of tumours ^10^, especially for ovarian cancers, where tumours are often detected only in late stages and are thus either spread in the abdomen or tumour mass is too big to allow for complete and unbiased sampling. Malignant ascites is reported in 40% of patients after neoadjuvant chemotherapy, making these patients candidates for genomic testing on cfDNA ^32^. cfDNA from ovarian cancer patients has been demonstrated to contain high tumour content and correlated well with tissue HRD scores ^10,32^.

Others have applied various strategies for the sampling of ascites fluid for the prediction of therapeutic outcome. Medina, et al. ^9^ suggested the use of a machine learning model integrating cfDNA fragmentomes and protein biomarkers that showed enhanced prognostic value compared to CA125 and HE4 (Human Epididymis Protein 4) alone. Werner, et al. ^10^ presented the feasibility, reliability and reproducibility for tracking tumour evolution with longitudinal sampling using targeted sequencing. Similar results have been obtained in our cohort. However, the retention of identified variants in cfDNA from ascites to *in vitro* patient avatars has not been quantified in those studies.

We have found good overlap between sampled ascites and ovarian cancer derived cell culture models. Variant retention was superior in organoids, which also showed an increase in heterozygous and homozygous variants. Similar findings were reported in PDAC organoids ^33^. On the other hand, spheroids retained other unique and low VAF variants highlighting the merits of each model.

Functional precision medicine is of growing importance for cancer therapy as a complement for genomic data ^11^. Here we could show the concordance in outcomes of the identification of functional variants via targeted sequencing with functional chemotherapeutic testing using the spheroid model. Importantly, while the culture of ascites-derived cells leads to the loss of unique variants, global resistance patterns match genomic data. These results have to be replicated by others using similar patient avatars in a controlled study setting such as in the OVAREX study ^34^.

This study underlines the suitability of cfDNA from ovarian cancer malignant ascites for mutational profiling supporting precision medicine. Ascites-derived cfDNA can provide clinically actionable information through a minimally invasive sampling method at key timepoints. Mutational profiling aligns well with cell culture models of the same patients using functional chemoresistance testing. These findings are particularly valuable at initial diagnosis, when ascites may precede interval debulking surgery, and at relapse, as an alternative to archived formalin-fixed tissue before adjuvant chemotherapy.

## Conclusions

Together, these results underline the suitability of cfDNA from ascites as a source for the identification of targetable driver variants that align well with functional chemoresistance testing in patient avatars. Like others, we therefore propose an implementation of ascites cfDNA sequencing to allow for precision medicine instead of invasive tissue biopsies. Additionally, drug response profiling of ascites derived cells in 3D-models enables a faithful reflection of tumor evolution and acquired resistance mechanisms leading to a better prediction of effective treatments, particularly in the situation of chemotherapy-resistant relapses.

## List of abbreviations

HGSOC: High-grade serous ovarian carcinomas
PFS: Progression-free survival
OS: Overall survival
cfDNA: cell-free DNA
OCM: ovarian cancer organoid medium
FT: fallopian tubes
ULA: ultra-low attachment
VAF: variant allele frequency
IC_50_: Half maximal inhibitory concentration
LGSOC: low-grade serous ovarian cancer
CUP: cancer of unknown progeny
CA125: Cancer Antigen 125/MUC16
HE4: Human Epididymal Protein 4

## Declarations

### Ethics approval and consent to participate

The studies involving humans were approved by the ethical committee of the Medical Faculty of the Christian-Albrechts-Universität zu Kiel (Ethics Approval 488/23 & 631/24). The studies were conducted in accordance with the local legislation and institutional requirements. The participants provided their written informed consent to participate in this study.

### Consent for publication

Not applicable.

### Availability of data and materials

All data generated or analysed during this study are included in this published article and its supplementary information files. The project is archived at the Competence Centre for Genomic Analysis (CCGA) Kiel under identifier 2994.

### Competing interests

QIAGEN was a subcontractor in the EU Horizon 2020 PCP project Instand-NGS4P, with the responsibility for developing an innovative diagnostic solution prototype for ovarian cancer. Employees of QIAGEN contributed to the preparation of this manuscript, and QIAGEN workflows and kits were used in parts of the experimental work. Franziska Kaiser, Kalle Günther, Katrin Rodenkirchen and Uwe Oelmüller are employees of QIAGEN GmbH Hilden, Germany. Kalle Günther, Katrin Rodenkirchen and Uwe Oelmüller participate in QIAGEN’s regular long term incentive program (LTIs). Uwe Oelmüller is Management Committee Co-Chair at PreAnalytiX (a QIAGEN/BD Company).

## Funding

The authors acknowledge funding from the University Hospital Schleswig-Holstein. DH, CR, & NH acknowledge funding by the University Cancer Center Schleswig-Holstein and University Cancer Center Hamburg via the Twinning Grant, CONNECT, and PHSH Shared-Campus-Awards Initiative. DH & NH thank UKSH-Gemeinsam Gutes tun! Förderstiftung des UKSH (017_2023) for funds enabling ovarian organoid cultures. DH and NH were further supported by funding from the German Research Society (DFG, ID 558466534) to NH. Sequencing work in this study was supported by Instand-NGS4P “INtegrated and Standardized NGS workflows for Personalized therapy”, co-funded by the European Union (Horizon 2020 grant no. 874719). The funders played no role in study design, data collection, analysis and interpretation of data, or the writing of this manuscript.

## Authors’ contributions

DH and NH supervised sample collection and processing. DH and AB performed cell culture experiments. DH, MR and MF performed bioinformatic analyses. CR, HDL, IA, NM, MTvM, and DOB collected samples and patient metadata. FK, KG, KR, UO, NH and JPW were decisive in panel design. MK and MB supported isolation of nucleic acids. AF and BB supported sequencing efforts. DH wrote the manuscript draft. NH conceptualized and supervised the study, selected patient samples, model systems and biomaterials. DB and NH established drug-panel design, NH established workflows for functional testing, DH performed and adapted functional testing and data-analysis. All authors read and approved the final manuscript.

## Acknowledgements

We thank Finja Grundt, Frauke Grohmann, and Catharina Verkooyen for their excellent technical support. We thank Dr. Reinhild Geisen, Dr. Anna Willms and the whole SYNENTEC team for the provision of the automated imaging system and technical assistance throughout live cell imaging. We sincerely thank Gudrun Kliem and Christian Klotz from the Robert Koch-Institute Berlin for quality control of the conditioned media.

